## Supplemental materials for "Structure-Function analysis of *Lactiplantibacillus plantarum* DltE reveals D-alanylated lipoteichoic acids as direct symbiotic cues supporting *Drosophila* juvenile growth"

### Extended Data

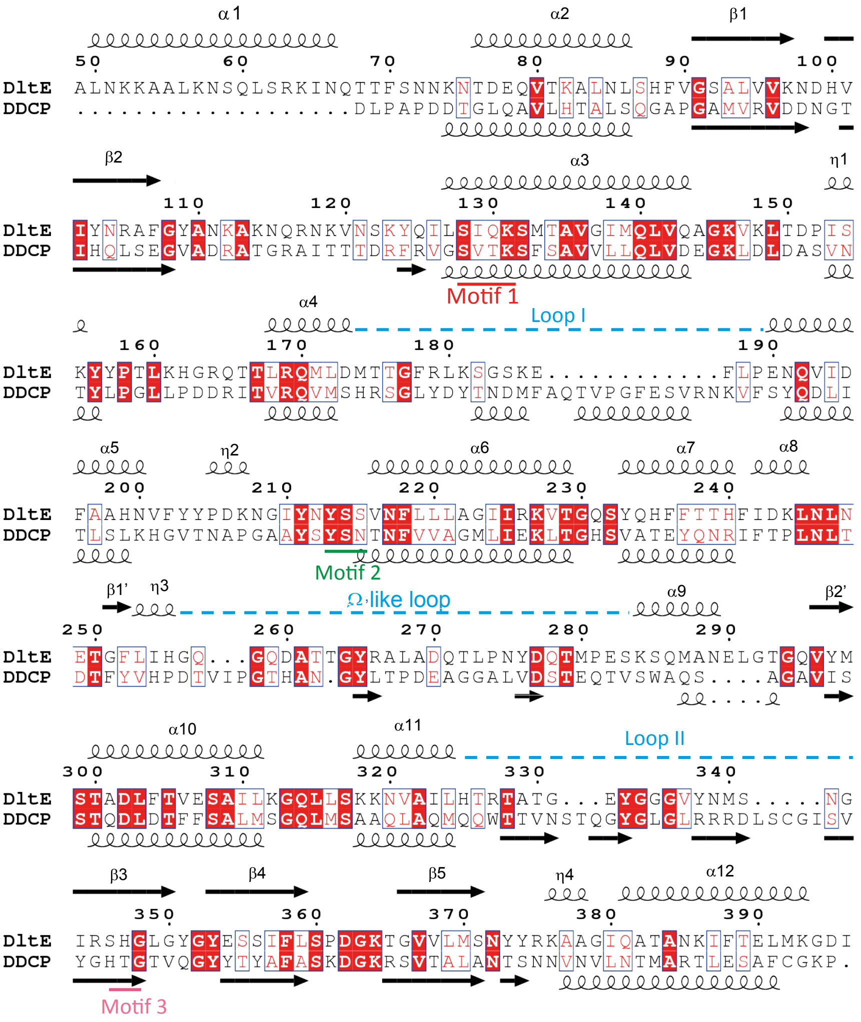


**Extended Data Fig. 1.** Sequence comparison of DltE with the *Streptomyces* R61 D-Ala-D-Ala carboxypeptidase (DDCP). Sequence alignment between DltE (residues 49-396) from *L. plantarum* and DDCP. The two proteins share a relatively low sequence identity of around 25%. The secondary structures extracted from the 3D X-ray structures are respectively depicted above (DltE, this study, PDB entry 8AJI) and below (DDCP, PDB entry 1HVB). Canonical serine type D-alanyl-D-alanine carboxypeptidases sequences exhibit three conserved motifs that are highlighted on the figure. The S-X-X-K motif 1(in red) encompasses a nucleophilic Ser which forms a catalytic dyad with the Lys residue. The S/Y-X-N motif 2 (in green) and the K/H-T/S-G motif 3 (in pink) complete the active site. Three loop regions that define the catalytic cavity architecture and differ between the two sequences are indicated on the alignment: Loop I, Loop II and the Ω-like loop. The figure was generated by ESPript (https://espript.ibcp.fr/).

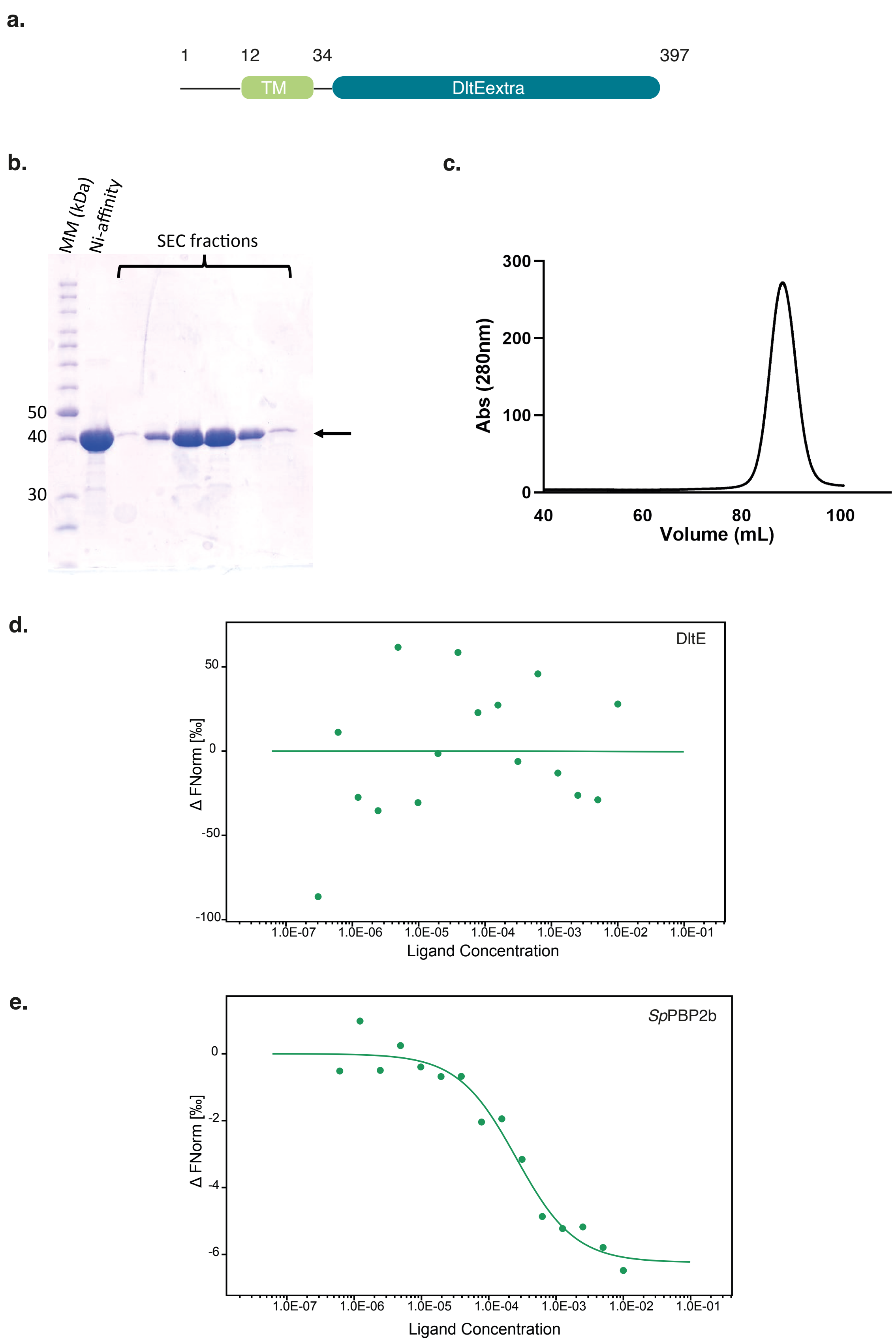


**Extended Data Fig. 2.** Production of DltE$_{extra}$ used for structure determination and biochemical assays. **a**, The *dltE* gene encodes for a 397 residues protein composed of a transmembrane segment predicted between residues 12 and 34 and a larger C-terminal extracellular region from residues 34 to 397 and named $DltE_{extra}$. **b**, SDS PAGE analysis of the purity of $DltE_{extra}$ after a two-step purification procedure including a Ni-Affinity and a size exclusion chromatography. The band corresponding to $DltE_{extra}$ (41kDa) is shown by an arrow. **c**, Size exclusion chromatography (SEC) and elution profile after injection of the Ni-affinity purified $DltE_{extra}$ on a Superdex 200 10/300 GL. **d**, **e**, MST normalized dose–response curves for the binding interaction between penicillin and $DltE_{extra}$ (**d**) and *S. pneumoniae* PBP2b used as a positive control (**e**) were obtained by plotting ΔFnorm against the ligand concentration. The data are representative of experiments made in triplicate. As expected from proteins from the PBP family *S. pneumoniae* PBP2 binds to penicillin while no binding was detected for DltE.

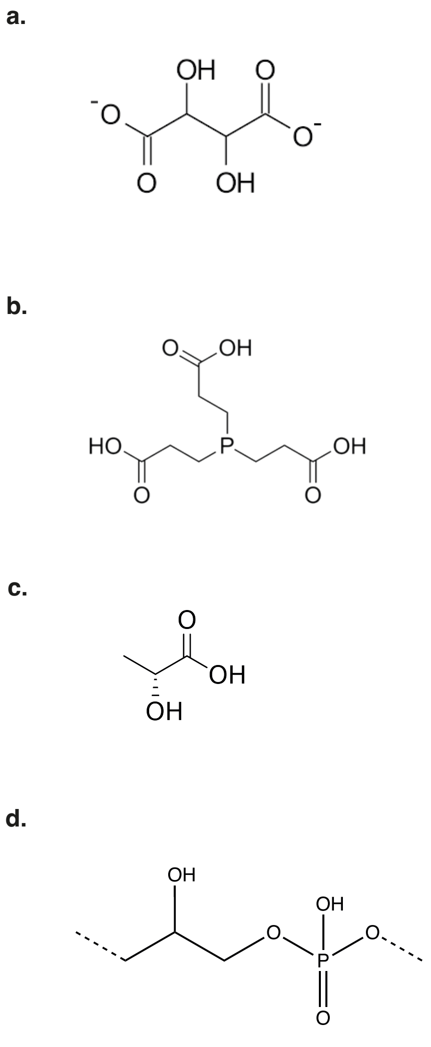


**Extended Data Fig. 3.** Chemical structures of **a**, Tartrate **b**, TCEP Tris(2-carboxyethyl)phosphine hydrochloride, **c**, D-Lac and **d**, subunit of *L. plantarum* LTA molecule without any substitution.

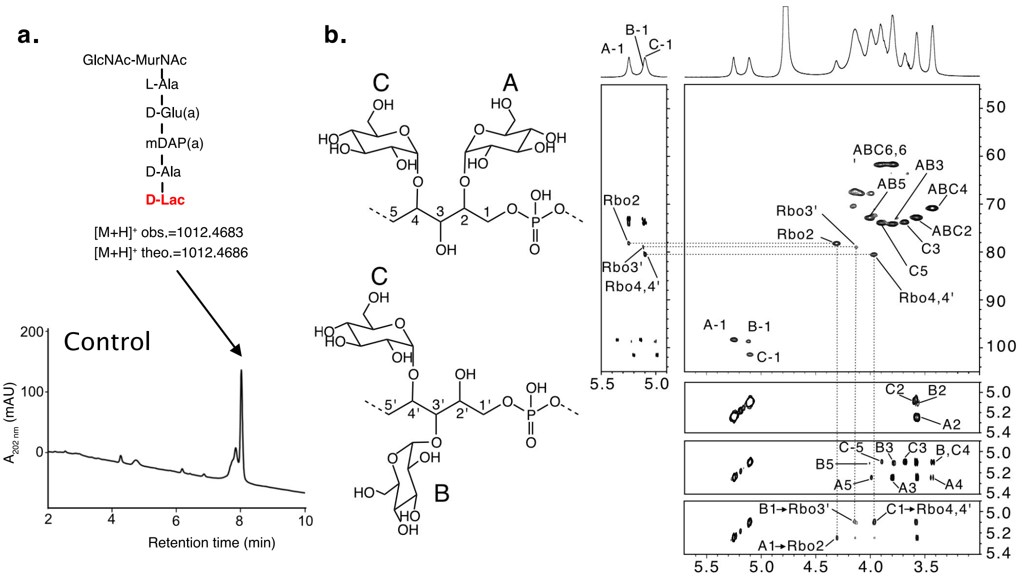


**Extended Data Fig. 4. a**, UHPLC analysis of the disaccharide-depsipentapeptide purified from the peptidoglycan of *Lp*$^{NC8}$ Δ*dacA1A2* mutant digested by mutanolysin. The muropeptide was identified by MS analysis. This muropeptide was used in activity tests of purified enzymes (DltE and DacA1) as shown in Fig. 3a. **b**, Multidimensional NMR analysis of WTA isolated from WT *L. plantarum* established the presence of two major repeating units made of phospho-ribitol (Rbo) differently substituted by Glc residues in C-2 position (residue A), C-3 position (residue B) and C-4 position (residue C) as shown on the left side. Individual spin systems of Glc residues A-C and Rbo positions 1-5 and 1'-5' were established from $^{1}$H-$^{13}$C HSQC (top left right), $^{1}$H-$^{1}$H COSY (second from top right) and $^{1}$H-$^{1}$H TOCSY (third from top right) spectra in agreement with literature that identified similar compounds from the cell wall of several strains of *L. plantarum*[1,2]. Linkage of αGlc residues A, B and C to positions 2, 3' and 4,4' of Rbo respectively were assigned from $^{1}$H-$^{1}$H NOESY (bottom, right) and $^{1}$H-$^{13}$C HMBC (top left) spectra as shown by the cross correlation signals.

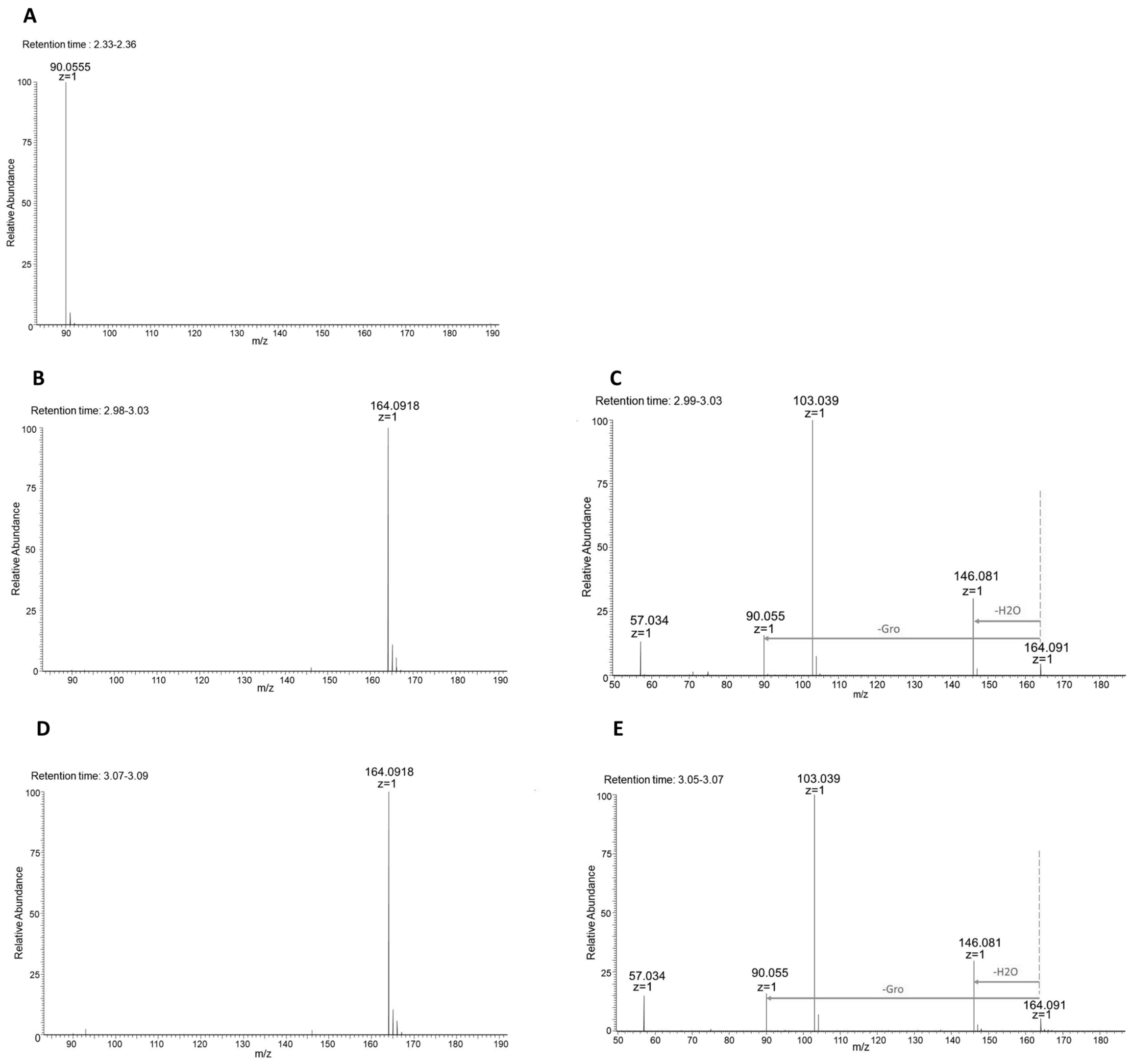


**Extended Data Fig. 5. a**, **b**, **d**, MS spectra of (**a**) Ala (retention time 2.33-2.36) (calculated [M+H]+ 90.0550), (**b**, **d**) Gro-Ala (retention time 2.98-3.03 and 3.07-3.09 (calculated [M+H]+ 164.0918). Corresponding chromatograms are shown in Fig. 3d. **c**, **e**, MS/MS spectra for peaks at retention time 2.98-3.03 and 3.07-3.09, confirming that both correspond to Gro-Ala.

**a.** **b.**

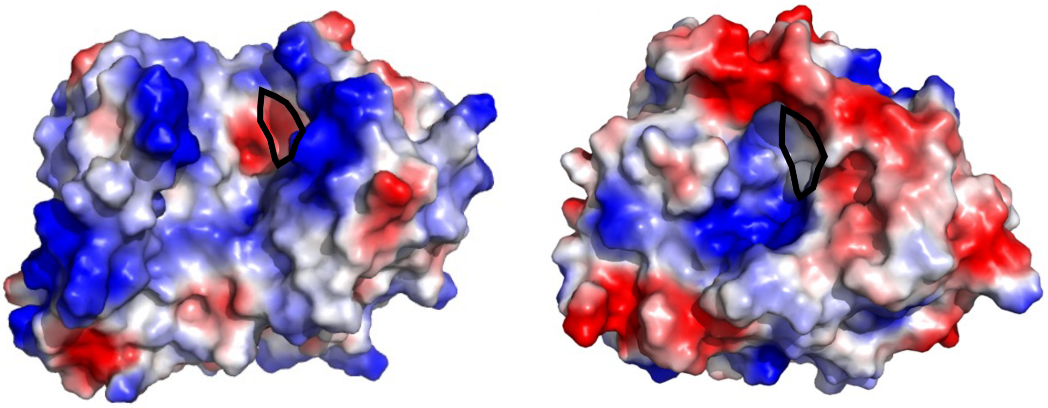

**Extended Data Fig. 6.** Molecular surfaces of DltE$_{extra}$ **a**, and DDCP **b**, colored according to the electrostatic potential. Residues are colored in blue and red according to positive and negative electrostatic potential, respectively. The apolar residues are colored white. The hydrophobic subsite suggested in DDCP to recognize the aliphatic portion of the peptide substrate is encircled in black (**b**). The corresponding region is marked as well in DltE (**a**) but is not conserved. It is replaced by mainly charged residues from the Ω-like loop and helix α9.

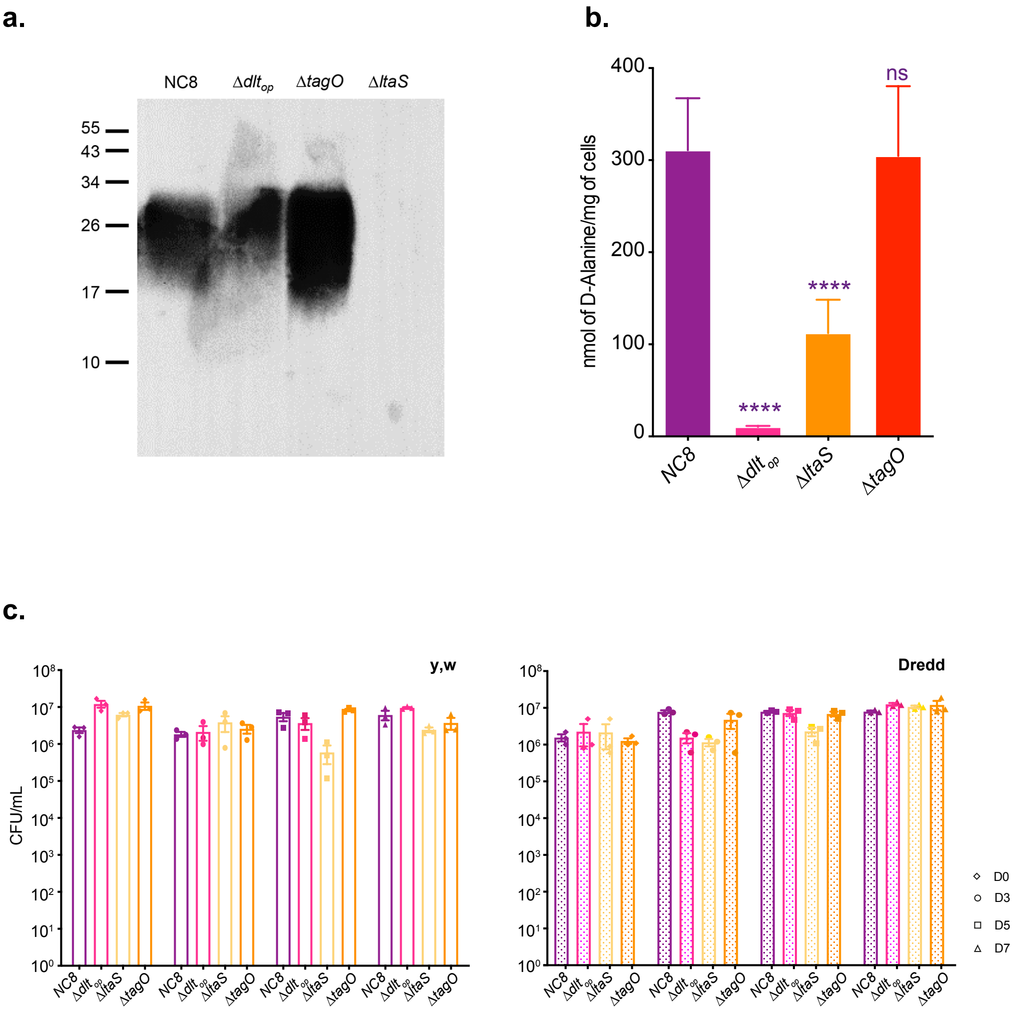


**Extended Data Fig. 7. a**, Western blot detection of LTA in wild-type $Lp^{NC8}$ and mutant derivatives. **b**, Amount of D-Ala released from whole cells of $Lp^{NC8}$, $\Delta dlt_{op}$, Δ*ltaS* and Δ*tagO* by alkaline hydrolysis and quantified by HPLC. Mean values were obtained from three independent cultures with two injections for each. Purple asterisks illustrate statistically significant difference with D-Ala released from $Lp^{NC8}$ . NS represents the absence of a statistically significant difference compared to $Lp^{NC8}$. ****: $P < 0.0001$. **c**, Evolution of the number of CFUs on fly food and larvae at days 3, 5, and 7 after inoculation with $Lp^{NC8}$, $\Delta dlt_{op}$, Δ*ltaS* and Δ*tagO* in *y,w* and *y,wDredd* flies.

### Supplementary Tables

**Supplementary Table 1.** Data collection and refinement statistics.

| | DltE $_{extra}$-apo | DltE $_{extra}$-tartrate | DltE $_{extra}$-TCEP | DltE $_{extra}$-TA |
|---|---|---|---|---|
| **Crystallization conditions** | 26% PEG MME 5K, 0.1M Tris pH8.5, 0.15M $LiSO_4$ and 30% ethylene glycol | 26% PEG 3350, 0.1M ammonium tartrate | 20% PEG 3350, 0.2M ammonium tartrate and 0.1M TCEP HCl | 26% PEG 3350, 0.1M ammonium tartrate<br><br>24h soaking with 10mM LTA |
| **Wavelength** | 0.87 | 0.97 | 0.87 | 0.98 |
| **Resolution range** | 47.75 - 1.86 (1.926 - 1.86) | 41.04 - 1.95 (2.02 - 1.95) | 49.45 - 1.94 (2.009 - 1.94) | 46.33 - 1.4 (1.45 - 1.4) |
| **Space group** | P 1 21 1 | P 21 21 21 | P 1 21 1 | P 21 21 21 |
| **Number of molecules in the a.u.** | 2[1] | 2[2] | 4[3] | 2[4] |
| **Unit cell (Å, °)** | 60.87, 95.712, 68.01, 90, 96.26, 90 | 48.69, 103.73, 152.53, 90, 90, 90 | 76.24, 98.64, 135.93, 90, 105.79, 90 | 48.68, 96.07, 150.83, 90, 90, 90 |
| **Total reflections** | 128187 (12709) | 113125 (11117) | 268566 (25402) | 279608 (27536) |
| **Unique reflections** | 64925 (6451) | 56870 (5568) | 139733 (13635) | 139817 (13770) |
| **Multiplicity** | 2.0 (2.0) | 2.0 (2.0) | 1.9 (1.9) | 2.0 (2.0) |
| **Completeness (%)** | 99.81 (99.92) | 99.32 (99.70) | 97.57 (95.26) | 99.98 (99.96) |
| **Mean I/sigma(I)** | 6.50 (2.16) | 15.08 (2.60) | 5.36 (2.49) | 16.61 (1.68) |
| **Wilson B-factor** | 14.15 | 32.35 | 21.70 | 15.31 |
| **R-merge** | 0.08491 (0.296) | 0.02214 (0.2728) | 0.06333 (0.2724) | 0.02281 (0.4422) |
| **R-meas** | 0.1201 (0.4186) | 0.03132 (0.3858) | 0.08957 (0.3853) | 0.03226 (0.6253) |
| **R-pim** | 0.08491 (0.296) | 0.02214 (0.2728) | 0.06333 (0.2724) | 0.02281 (0.4422) |
| **CC1/2** | 0.988 (0.791) | 0.999 (0.852) | 0.99 (0.822) | 1 (0.678) |
| **CC*** | 0.997 (0.94) | 1 (0.959) | 0.997 (0.95) | 1 (0.899) |
| **Reflections used in refinement** | 64911 (6451) | 56861 (5567) | 139689 (13635) | 139809 (13766) |
| **Reflections used for R-free** | 3142 (288) | 2810 (293) | 1993 (186) | 1053 (103) |

| | | | | |
|---|---|---|---|---|
| **R-work** | 0.1715 (0.2609) | 0.1877 (0.3129) | 0.1869 (0.2420) | 0.1742 (0.2679) |
| **R-free** | 0.2072 (0.3159) | 0.2206 (0.3461) | 0.2105 (0.2730) | 0.1929 (0.3072) |
| **CC(work)** | 0.959 (0.879) | 0.956 (0.819) | 0.952 (0.798) | 0.965 (0.811) |
| **CC(free)** | 0.949 (0.845) | 0.959 (0.750) | 0.925 (0.806) | 0.968 (0.784) |
| **Number of non-hydrogen atoms** | 6008 | 5656 | 11962 | 5681 |
| **macromolecules** | 5132 | 5054 | 10610 | 5070 |
| **ligands** | 31 | 30 | 142 | 109 |
| **solvent** | 845 | 572 | 1210 | 534 |
| **Protein residues** | 647 | 642 | 1347 | 641 |
| **RMS(bonds)** | 0.010 | 0.009 | 0.010 | 0.009 |
| **RMS(angles)** | 1.34 | 1.52 | 1.38 | 1.11 |
| **Ramachandran favored (%)** | 97.98 | 97.49 | 96.56 | 97.33 |
| **Ramachandran allowed (%)** | 1.71 | 2.19 | 2.91 | 2.35 |
| **Ramachandran outliers (%)** | 0.31 | 0.31 | 0.52 | 0.31 |
| **Rotamer outliers (%)** | 0.18 | 2.39 | 1.22 | 0.18 |
| **Clashscore** | 2.03 | 2.66 | 3.53 | 3.79 |
| **Average B-factor** | 17.43 | 35.01 | 28.11 | 19.83 |
| **macromolecules** | 15.66 | 34.08 | 27.08 | 18.61 |
| **ligands** | 26.46 | 60.20 | 54.31 | 33.34 |
| **solvent** | 27.89 | 41.96 | 34.11 | 29.45 |

Statistics for the highest-resolution shell are shown in parentheses.

[1]The final model contains residues 72-393 in chain A and residues 69-393 in chain B

[2]The final model contains residues 75-394 in chain A and residues 72-393 in in chain B

[3]The final model contains residues 49-396 in chain A, residues 70-393 in chain B, residues 64-394 in chain C and residues 53-396 in chain D. This structure is the most complete.

[4]The final model contains residues 75-395 in chain A and residues 75-394 in in chain B

**Supplementary Table 2.** Proton and carbon chemical shifts of Glc, Gro and Ala constituents of LTA purified from *L. plantarum* WT.

| | | Chemical Shifts (ppm) | | | | | | |
|---|---|---|---|---|---|---|---|---|
| | | 1 | 2 | 3 | 4 | 5 | 6a | 6b |
| **Gro A** | H (δ) | 3.90 - 3.96 | 4.06 | 3.90 - 3.96 | | | | |
| | C (δ) | 64.7 | 70.6 | 64.7 | | | | |
| **Gro B** | H (δ) | 4.11 | 5.40 | 4.11 | | | | |
| | C (δ) | 64.7 | 75.3 | 64.7 | | | | |
| **Gro C,D** | H (δ) | 4.02 | 4.13 | 4.02 | | | | |
| | C (δ) | 66.0 | 76.4 | 66.0 | | | | |
| **αGlc C** | H (δ) | 5.19 | 3.55 | 3.77 | 3.42 | 3.93 | 3.77 | 3.88 |
| | C (δ) | 98.9 | 72.6 | 74.0 | 70.8 | 73.1 | 61.56 | |
| **αGlc D** | H (δ) | 5.19 | 3.55 | 3.77 | 3.42 | 4.18 | 4.45 | 4.63 |
| | C (δ) | 98.9 | 72.6 | 74.0 | 70.8 | 70.9 | 66.2 | |
| **Ala** | H (δ) | | 4.29 | 1.63 | | | | |
| | C (δ) | 172.5 | 49.9 | 16.36 | | | | |

**Supplementary Table 3.** Primers used in this study.

| Primer name | Sequence (5'→3')* | Reference |
|---|---|---|
| XL01 | CTTGATATCGAATTCCTGCACTTGATTCAAAATCAAGAGACCCT | This study |
| XL02 | TACCATGCCTGATTAATCGAACTCGTATCAACTAAGGG | This study |
| XL03 | TCGATTAATCAGGCATGGTAATTTCTTCCTCCG | This study |
| XL04 | AGTGGATCCCCCGGGCTGCAACGTGCTCAGGCGTGTTGAA | This study |
| XL05 | CTTGATATCGAATTCCTGCAGACACCGGCATCCTTATTAA | This study |
| XL06 | CACAAATGATCCATTAAAAACCAAATAAATCATTGA | This study |
| XL07 | GGTTTTTAATGGATCATTTGTGCGTAACTCCCT | This study |
| XL08 | AGTGGATCCCCCGGGCTGCACCGAATCCACGTGCACTATA | This study |
| XL09 | AGTGGATCCCCCGGGCTGCAAACAGTACCAATCAGAAGAGGA | This study |
| XL10 | CTATAATTTAGTTTCTCATTTCTCAATTATCCCTTTCT | This study |
| XL11 | TTGAGAAATGAGAAACTAAATTATAGCAGTTAGTGT | This study |
| XL12 | CTTGATATCGAATTCCTGCAGTAACTGGTTTAAGATCAGCCGT | This study |
| XL13 | AGTGGATCCCCCGGGCTGCAGCCATGTTAATTGGTTTTCA | This study |
| XL14 | GAGCATCACTTTAACATAGTACCTTCCTTTAATTCGT | This study |
| XL15 | GTACTATGTTAAAGTGATGCTCGCTTAATAGATCG | This study |
| XL16 | CTTGATATCGAATTCCTGCATACGGTAGCGACCACGTCTC | This study |
| XL17 | TGGATCCCCCGGGCTGCAATGCGGCTTCAAAATCAAGGT | This study |
| XL18 | ATCGATAAAATTTGATACTTTGAATTGACTTTATTACGT | This study |
| XL19 | CAAAGTATCAAATTTTAGCAATTCAAAAGTCAATGACGG | This study |
| XL20 | TGATATCGAATTCCTGCACTGGTCAGGCAATCCGAAGT | This study |
| rp49f | GACGCTTCAAGGGACAGTATCTG | [3] |
| rp49r | AAACGCGGTTCTGCATGA | [3] |
| jon66ciif | AAACTGACCCCGGTCCAC | [3] |
| jon66ciir | CCTCCCAG CCGAT AGC | [3] |
| jon65Aif | CAACAACTACCAGGCTGGTG | [3] |
| jon65Air | GCCCTCATCGGAGGTCTT | [3] |

*Overlapping sequences for Gibson assembly are underlined.

**Supplementary Table 4.** Bacterial strains and plasmids used in this study.

| Strain or plasmid | Relevant characteristics | Reference or source |
|---|---|---|
| **Strain** | | |
| ***E. coli*** | | |
| TG1 | *supE hsd5h thi* ($\Delta$*lac-proAB) F' (traD36 proAB-lacZ*$\Delta$*M15)* | [4] |
| GM1674 | $dam^{-} dcm^{-}$ $repA^{+}$ | [5] |
| ***L. plantarum*** | | |
| NC8 | Isolated from grass silage, plasmid free | [6] |
| $\Delta$*dltE* | NC8 strain deleted for *nc8_1738* (formerly *pbpX2*) | [7] |
| $\Delta$*dltXABCD* | NC8 strain deleted from *nc8_1737* to *nc8_1733* | [7] |
| $\Delta dlt_{op}$ | NC8 strain deleted from *nc8_1738* to *nc8_1733* | [7] |
| $\Delta$*ltaS* | NC8 strain deleted for *nc8_1125* | This study |
| $\Delta$*tagO* | NC8 strain deleted for *nc8_0646* | This study |
| $\Delta$*dacA1A2* | NC8 strain deleted for *nc8_2720* and *nc8_1122* | This study |
| $dltE^{S128A}$ | Knock-in of *dltE* mutated on S128 on $\Delta$*dltE* strain | This study |
| **Plasmids** | | |
| pG+host9 | $Erm^{r}$, repATs | [8] |

**Supplementary Table 5.** Primers used for *E. coli* plasmid constructions.

| Number | Name | Sequence (5' to 3') | Reference |
|---|---|---|---|
| *1* | 5-pPbpX2extra | AGATATACCATGGCTACAGAGCGGCAAGCAGC | This study |
| *2* | 3-pPbpX2 extra | TCGACTCGAGCTTAATATCACCCTTCATTAATTCCGTAAAAATCTTG | This study |
| *3* | 5-pPbpX2 extra S128A | GTTGCTAGCTGGCATTATTCGC | This study |
| *4* | 3-pPbpX2 extra S128A | GCGAATAATGCCAGCTAGCAAC | This study |

**Supplementary Table 6.** Plasmids used in this study

| Plasmid | Description and main characteristics | Source | Primers |
|---|---|---|---|
| pET-28a(+) | T7 promoter, C-terminal 6×His, $Kan^R$ | Novagen | - |
| pPbpX2 $_{extra}$ | pET-28a(+) derivative encoding *L. plantarum* wild-type PbpX2 extracellular domain (residues 34-397) fused to a C-terminal $(His)_6$ tag | This study | 1,2 |
| pPbpX2 $_{extra}$ S128A | pET-28a(+) derivative encoding *L. plantarum* PbpX2 extracellular domain S128A mutant fused to a C-terminal $(His)_6$ tag | This study | 3,4 |
